# Assisted assembly of bacteriophage T7 core components for genome translocation across the bacterial envelope

**DOI:** 10.1101/2021.01.04.424714

**Authors:** Mar Pérez-Ruiz, Mar Pulido-Cid, Juan Román Luque-Ortega, José María Valpuesta, Ana Cuervo, José L. Carrascosa

## Abstract

In most bacteriophages, the genome transport across bacterial envelopes is carried out by the tail machinery. In *Podoviridae* viruses, where the tail is not long enough to traverse the bacterial wall, it has been postulated that viral core proteins are translocated and assembled into a tube within the periplasm. T7 bacteriophage, a member from the *Podoviridae* family, infects *E. coli* gram-negative bacteria. Despite extensive studies, the precise mechanism by which this virus translocates its genome remains unknown. Using cryo-electron microscopy, we have resolved the structure two different assemblies of the T7 bacteriophage DNA translocation complex, built by core proteins gp15 and gp16. Gp15 alone forms a partially folded hexamer, which is further assembled by interaction with gp16, resulting in a tubular structure with dimensions compatible with traversing the bacterial envelope and a channel that allows DNA passage. The structure of the gp15-gp16 complex also shows the location in gp16 of a canonical transglycosylase motif essential in the bacterial peptidoglycan layer degradation. Altogether these results allow us to propose a model for the assembly of the core translocation complex in the periplasm, which helps in the understanding at the molecular level of the mechanism involved in the T7 viral DNA release in the bacterial cytoplasm.

**SIGNIFICANCE STATEMENT:** T7 bacteriophage infects *E. coli* bacteria. During this process, the DNA transverses the bacterial cell wall, but the precise mechanism used by the virus remains unknown. Previous studies suggested that proteins found inside the viral capsid (core proteins) disassemble and reassemble in the bacterial periplasm to form a DNA translocation channel. In this article we solved by cryo-electron microscopy two different assemblies of the core proteins that reveal the steps followed by them to finally form a tube large enough to traverse the periplasm, as well as the location of the transglycosylase enzyme involved in peptidoglycan degradation. These findings confirm previously postulated hypothesis and make experimentally visible the mechanism of DNA transport trough the bacterial wall.

## INTRODUCTION

Bacteriophages (phages) are viruses that infect bacteria, and are considered to be the most abundant entities on Earth. Caudovirales are double-stranded DNA (dsDNA) tailed phages (1, 2), which can infect a wide variety of hosts, and are present in many different environments (3). These viruses are composed of an icosahedral capsid with a tail protein complex, which is assembled at one special vertex of the capsid, the portal vertex (4). The connector or portal protein builds an entry channel to the viral capsid and serves as a docking point for the tail complex (4, 5). Tailed phages generally show a common infection strategy and thus have strong structural homologies, although the protein machinery responsible for host adsorption has specific characteristics depending on each phage family (3). Some phages such as the *Podoviridae* present a short, non-contractile tail that is not long enough to traverse the thick bacterial wall (6), and some others do not present tail machinery at all. These viruses have developed an alternative mechanism to cross the bacterial envelope using internal capsid proteins or membranes that are able to assemble tubular structures during infection, thus directing DNA translocation through the bacterial wall (7–11).

Bacteriophage T7 is a well-characterized *E. coli* virus member of the *Podoviridae* family. The viral particle is composed of a 55-nm icosahedral capsid and a 23-nm short non-contractile tail (12). The most remarkable structure of the T7 viral particle is the internal core, a cylindrical structure that is ~290 Å long and ~170 Å wide located on top of the connector, which stabilizes the packaged DNA inside the capsid (13). This complex is made up of three proteins: gp14 (~20.8 kDa), gp15 (~84.2 kDa) and gp16 (~144 kDa) (9, 13). These internal core proteins are not essential for morphogenesis of the viral capsid, but they are required to translocate the viral genome during the T7 infection process (11, 14, 15). A lytic transglycosylase motif present in gp16 is essential during infection to overcome the highly cross-linked peptidoglycan (16–18). As it is the case for most phages, T7 first binds reversibly to a primary receptor that allows the correct tail orientation related with the bacterial surface. Then, an irreversible interaction takes place with the rough lipopolysaccharide (LPS) (19), which causes conformational changes in the tail leading to the opening of the channel. Later a tubular conduit, probably built by the core proteins, is assembled. It crosses the bacterial wall (9, 11), although it is not clear how this is accomplished. One hypothesis was proposed by Hu *et al.* (11) in a study using cryo-electron tomography, where they described the presence of transient tubular structures during T7 infection. They proposed that the core complex formed by gp14, gp15 and gp16 could disassemble after adsorption and pass through the open tail channel in a completely (or partially) unfolded state (11). According to this hypothesis, partially unfolded gp14 should be ejected through the channel of the tail complex, and then refolded to form a pore into the outer membrane (OM) of *E. coli*, which allows passage of unfolded gp15 and gp16 to cross (11, 18). Once in the periplasm, gp15 and gp16 oligomerize as a tubular structure that spans the periplasm and internal membrane (IM) and reaches the cytoplasm. When the channel formed by gp14, gp15 and gp16 is complete, translocation of the T7 genome into the bacteria cytoplasm can take place.

To date no atomic structures of the T7 bacteriophage core components gp14, gp15 or gp16 have been solved. Here, we report two cryo-electron microscopy (cryo-EM) structures of the bacteriophage T7 core assemblies: gp15 alone (505 kDa), and that of the complex formed by gp15 and gp16 (gp15-gp16 complex; 1.365 MDa). The gp15 protein alone forms a tubular structure *in vitro*, although its C-terminal half part is disordered. The complex formed by gp15 and gp16 is also tubular structure, but this time with a fully folded gp15. Although only a small fragment of gp16 is observed in the structure of the complex, the solved model comprises the transglycosylase domain. This is the first time that it has been possible to show that these bacteriophage proteins, which form part in the mature virus of the fully structured core complex, are able during the infection process to unfold to exit the phage and reassemble into a tubular structure of a size compatible with the translocation of viral DNA across the bacterial envelope.

## RESULTS

### Phage T7 gp15 forms tubular oligomers *in vitro*

Gp15 recombinantly expressed and purified as described in the Methods section (Supplementary Fig. 1A), was subsequently characterized by analytical centrifugation (AUC) (Supplementary Fig. 1D) and dynamic light scattering (DLS) (Supplementary Fig. 1E). The two techniques revealed that more than 80% of the sample was present as an oligomer of ~ 540 kDa, compatible with the presence of an hexamer, with less than 10% as a monomer, regardless of the concentration of sample used (from 0.25 mg/ml to 1 mg/ml) (Supplementary Fig. 1 D and E). Aliquots of the purified protein were negatively stained and observed by transmission electron microscopy (Supplementary Fig. 2A). A total of 1648 particles were selected and processed as described in the Methods section. The 2D classification rendered two main classes (Supplementary Fig. 2B), one with a cylindrical shape and hosting a channel, and the other, presumably the orthogonal view, with a six-fold symmetry that strengthens the notion of gp15 forming an hexamer. The 3D reconstruction confirms the tubular shape of the oligomer, of ~ 130 Å long and an average width of ~ 100 Å (Supplementary Fig. 2C).

We next resorted to cryo-EM to try to generate a high-resolution structure of the gp15 oligomer. For that, aliquots of gp15 were vitrified as described in the Methods section. A FEI-Thermofischer Talos Arctica equipped with a Falcon III was used for data acquisition. A total of 2470 movies were acquired (Supplementary Fig. 3A) and 444.807 total particles were picked from which, after classification 50.980 were selected as described in the Methods section (Supplementary Fig. 3B). A 3.64 Å resolution map of the oligomer was obtained in which C6 symmetry was applied (Fig. 1 and Supplementary Fig. 3C-E). The resolution seems to be very similar throughout all the 3D reconstruction, which points to a very rigid structure being formed (Supplementary Fig. 3E). The shape of the oligomeric gp15 is that of a nozzle of 140 Å length and 88 Å at the widest point (Fig. 1A). A channel runs along the longitudinal axis, with a wider entrance at one end (36 Å) and a narrower one at the other (23 Å), but always wide enough to allow the passage of DNA (Fig. 1B).

**Figure 1.**
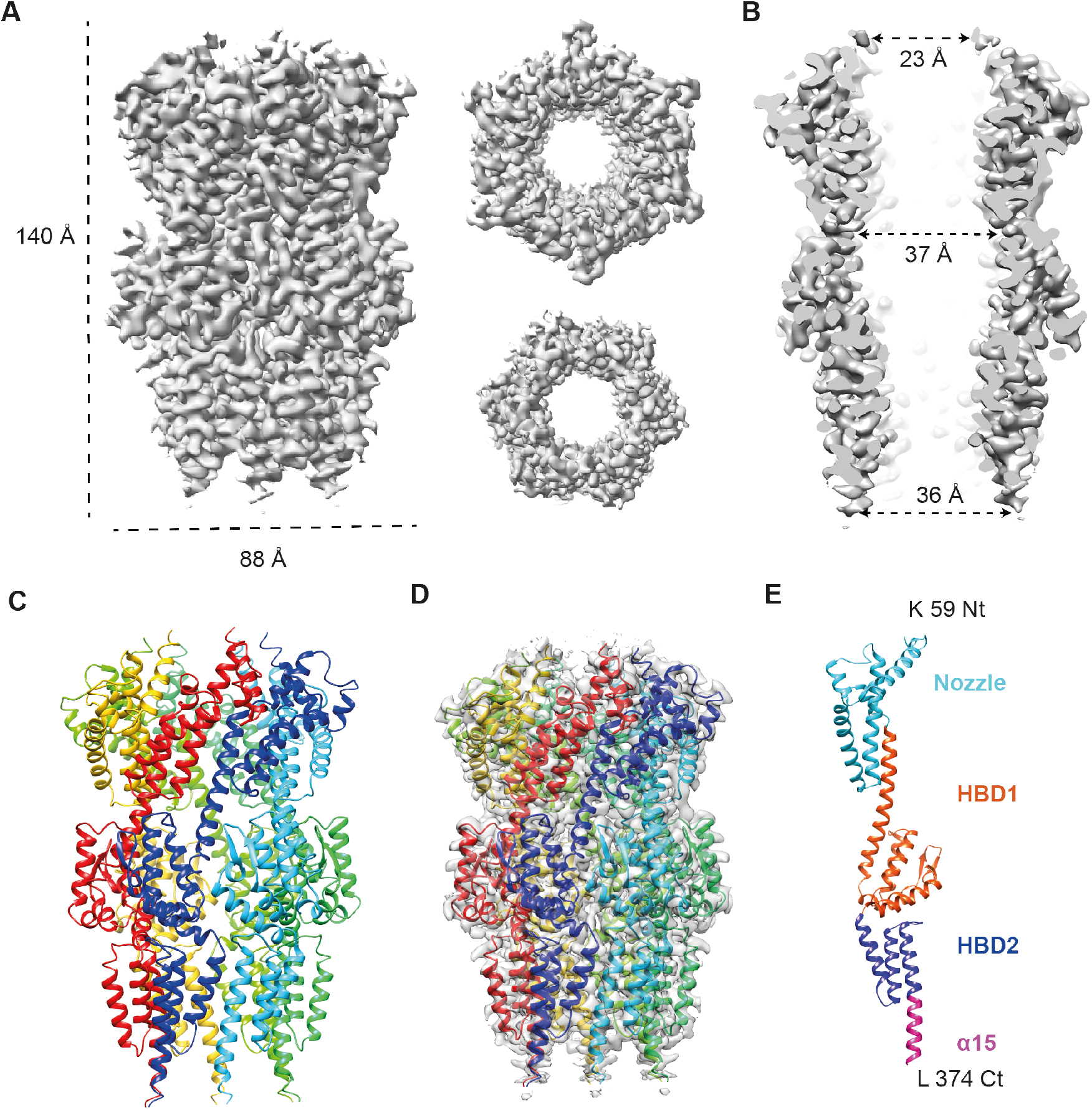
Structure of the bacteriophage T7 gp15 core protein. **A.** Side, top and bottom views of the cryoEM map of gp15 assembly at 3.6 Å resolution showing the dimensions of the complex. **B.** Longitudinal cut of gp15 cryoEM density map showing the dimensions of the internal channel. **C.** Ribbon representation of the gp15 atomic model. Each monomer is depicted in a different color. **D.** Docking of the gp15 atomic model into the corresponding cryoEM density map. **E.** Ribbon representation of a gp15 monomer; the structure is colored showing the different domains, and the N- and C-terminal residues are labeled.

The high resolution of the map allowed us an *ab initio* tracing of the gp15 polypeptide chain, from residues K59 to L374, that encompassed 15 α-helices and 2 β-strands (Fig. 1C-E and Supplementary Fig. 3F). The structure is composed of three helical bundle domains (HBD) that we have named nozzle (residues 59-170), HBD1 (residues 171-302) and HBD2, which contains the long α-helix 15 (residues 303-374) (Fig. 1E). This leaves undetermined the first 58 residues and the last 373 ones. The absence of more than half of the protein density in the reconstructed hexamer could be due either to protein degradation, domain flexibility or partial folding. However, our analytical ultracentrifugation and dynamic light scattering (DLS) assays (Supplementary Fig. 1 D and E) unambiguously confirmed the presence of an oligomer whose molecular mass is that of a full-length hexameric gp15. The possibility of the oligomer being fragmented during the vitrification process (20) could also be ruled out as the dimensions of the gp15 oligomer 3D reconstructed using negative staining agree with those of the vitrified complex (compare Fig. 1A with Supplementary Fig. 2C). However, it is interesting to note a blurry area in the 2D classes of both negatively-stained and vitrified samples, in the area that we assigned to the C-terminal region (arrow in Supplementary Figs. 2B and 3B). This suggests that this region of the polypeptide chain is disordered rather than degraded. In summary, purified gp15 uses its N-terminal half to form a tubular, hexameric structure, whereas the C-terminal half is disordered and does not contribute to the final structure.

### T7 gp16 acts as a chaperone and promotes the folding of the full structure of the gp15 hexamer

The gp16 core protein was also recombinantly expressed and purified (Supplementaty Fig. 1B), with a monomeric mass of ~ 150 kDa, similar to what it is predicted from its aminoacid sequence. The gp16 oligomeric state was analyzed using analytical ultracentrifugation and DLS and the two techniques pointed to the protein being present as a monomer (Supplementary Fig. 1D and F), and this was confirmed by negative-staining electron microscopy, which only revealed the presence of proteins of small size and aggregates (Supplementary Fig. 2D), and prevented the use of cryo-EM for its higher-resolution characterization.

Since the internal core of bacteriophage T7 is formed by proteins gp14, gp15 and gp16, we next wanted to know what sort of structure is formed by proteins gp15 and gp16 and for that we coincubated both proteins, which resulted in the formation of a high-molecular mass complex. AUC experiments revealed, together with the presence of free gp16, another peak with an experimental sedimentation coefficient of 15.1 S, a value whose correction to standard conditions (s20, w = 23.6 S) was compatible with a molecular mass of around 1.4 MDa (Supplementary Fig. 1D). This molecular mass is in agreement with the presence of a complex made of the hexameric gp15 already established (500 kDa) and another six monomers of gp16 (~ 865 kDa).

The gp15-gp16 complex was next purified by ultracentrifugation in a glycerol gradient, which confirmed the formation of the high-molecular mass complex (compare the fraction of the gp15-gp16 complex with regard to those of gp15 or gp16; Supplementary Fig. 1C). The complex was first analyzed by negative-staining EM that showed and homogeneous tubular assembly of similar shape to that of gp15, albeit much longer (Supplementary Fig. 2E). This was confirmed by the 3D reconstruction of the gp15-gp16 complex. A total of 2553 particles were used for a 2D classification, which again revealed two main classes (Supplementary Fig. 2F), one corresponding to the end-on view, which clearly showed the six-fold symmetry already observed in the gp15 oligomer, and a side view with a cylindrical shape, again similar, albeit longer to that of gp15. The 3D reconstruction carried out using these classes and imposing six-fold symmetry reinforces the notion of the gp15-gp16 complex having a tubular structure of ~ 250 Å long and an average width of ~ 120 Å (Supplementary Fig. 2G).

We next resorted to cryo-EM to generate a high-resolution structure of the gp15-gp16 complex. For that, aliquots of gp15-gp16 were vitrified and the best grids were loaded onto a FEI-Thermo fisher Titan Krios cryoelectron microscope operated at 300 kV (at the eBIC, Diamond Light Source, UK), equipped with a Gatan K2 summit detector. A total of 5,506 movies were acquired (Supplementary Fig. 4A) and 828.762 total particles were picked from which after classification 72.882 were selected as described in the Methods section (Supplementary Fig. 4B). A 3.19 Å resolution map of the gp15-gp16 complex with imposed C6 symmetry was finally generated (Fig. 2 and Supplementary Fig. 4C-E). The 3D reconstruction of the complex reveals a tubular structure of 210 Å long, 70 Å longer than the gp15 oligomer, and 138 Å at its widest point (Fig. 2A). The tube is an asymmetric one, with two different halves, one narrower and identical to the gp15 oligomeric structure (Fig. 2B), and a wider one with two protrusions, one roughly at the center of the tube that we called wing (arrow in Fig. 2A) and the other at its widest point (Fig. 2A and B). The channel that runs along the whole structure is widest in its central part (31-37 Å) and has the two narrowest points at the two entrances of the structure (21 and 23 Å; Fig. 2C).

**Figure 2.**
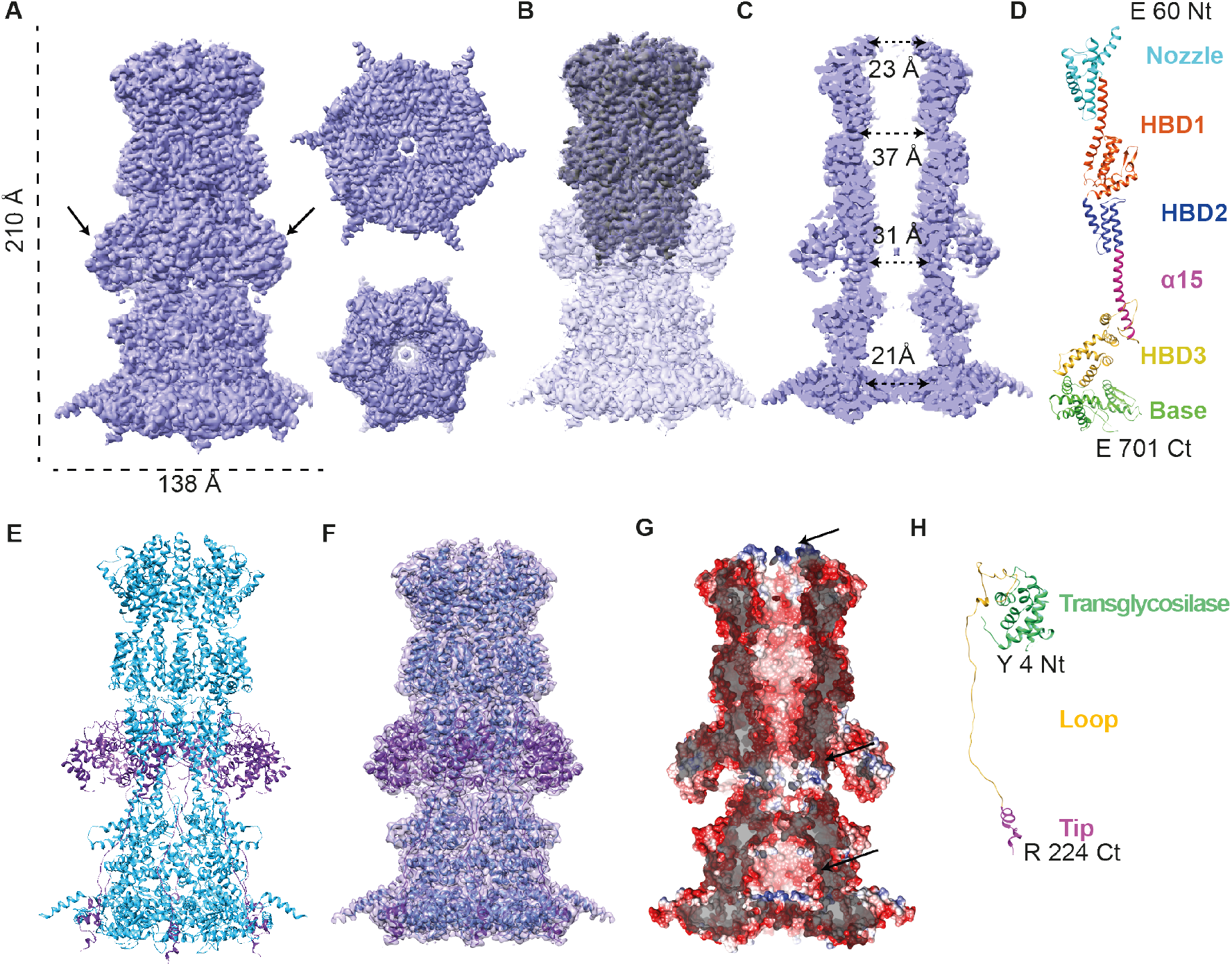
Structure of the bacteriophage T7 gp15-gp16 core complex. **A.** Side view, top and bottom views of the cryoEM map of gp15-gp16 core complex at 3.19 Å resolution showing the dimensions of the complex. The wing domain is pointed with an arrow. **B.** Overlapping of the gp15-gp16 complex (transparent) and gp15 cryoEM maps (solid). **C.** Longitudinal cut showing the dimensions of the internal channel. **D.** Ribbon representation of gp15, the structure is colored showing the different domains, and the N- and C-terminal residues are labeled. **E.** Ribbon representation of the gp15-gp16 atomic model, gp15 is shown in turquoise and gp16 in purple. **F.** Overlapping of the gp15-gp16 atomic model with its cryoEM density map. **G**. Electrostatic potential showing a longitudinal cut of gp15-gp16 indicating the inner channel surface (right). Blue color represents a 10 kcal / (mol · *e*) positive potential, while red represents a −10 kcal / (mol· *e*) negative potential. The electropositive charged rings in the internal channel are highlighted with arrows**. H.** Ribbon representation gp16 monomer, the structure is colored showing the different domains, and the N- and C-terminal residues are labeled.

As with the gp15 oligomer, *ab initio* tracing of the polypeptide chain of the gp15-gp16 complex was straightforward (Fig. 2D-F) except for the wings, which were less defined (Supplementary Fig. 4D and E; see below). As described above, one half of the structure coincided with that of the gp15 hexamer and the tracing of this part gave identical results (Fig. 2B, E and F). Surprisingly, tracing of the rest of the density showed that the majority of the newly reconstructed polypeptide chain corresponded to the regions of the gp15 peptide that were not visible in the 3D reconstruction of the gp15 oligomer. This time we were able trace the gp15 polypeptide chain from residues E60 to E701, which included 30 α-helices and 4 β-strands (Fig. 2D-F and Supplementary Fig. 4G), and that now can be divided in 6 domains: the previously described nozzle domain, HBD1 and HBD2 plus the newly visualized α-15 (now from residues A303-N396), HBD3 (residues G397-N524) and base domain (residues G525-E701) (Fig. 2D). The 3D reconstruction reveals that gp15 builds entirely the channel of the core complex, and the analysis of the electrostatic surface of the channel walls show an overall negative charge with three positive rings at the nozzle, α15 and basal domains (Fig. 2G). The atomic structure of the gp15 in complex with gp16 strongly suggests that our first interpretation for the gp15 hexamer structure was correct, and the regions not previously observed in the gp15 hexamer were indeed disordered. This clearly points to gp16 as a chaperone that assists gp15 in reaching its full, native structure.

### T7 gp16 shows a conserved transglycosylase active site that is visualized in the 3D reconstruction of the gp15-gp16 complex

As the gp15 sequence was almost completely traced (Supplementary Fig. 4G), the only regions not determined in the volume of the gp15-gp16 complex were the wings, which were assigned to gp16 (Fig. 2E and F). Although the wings presented lower resolution than the rest of the structure (Supplementary Fig. 4E), the density was good enough as to allow the tracing of 196 residues. The structure determined, when compared to others in the DALI server (21), had a high similarity to that of the *E. coli* SltY lytic transglycosilase (Fig. 3). This agreed with previous studies (17, 22) that had already shown that the N-terminal sequence of the T7 gp16 protein presents homology with the C-terminal sequence of the *E. coli* lytic transglycosylase SltY (Fig. 3), and this reinforced the tracing of a part of gp16, which encompasses residues Y4 to R224 with the exception of two loops (residues F40-M54 and L66-R76). The reconstructed part of gp16 can be divided in three parts: the transglycosylase domain (residues Y4-V145), a long disordered region, the loop domain (residues A146-P205) and two small helices that we called the tip domain (residues F206-R224) (Fig. 2H). These results show that the gp15 and gp16 interact to form a stable complex in which gp15, which by itself is an oligomer with only half of its sequence structured, gets completely folded with the assistance of gp16. Paradoxically, although gp16 acts as a gp15 chaperone, when binding to gp15 maintains disordered most of its sequence (~ 85%).

**Figure 3.**
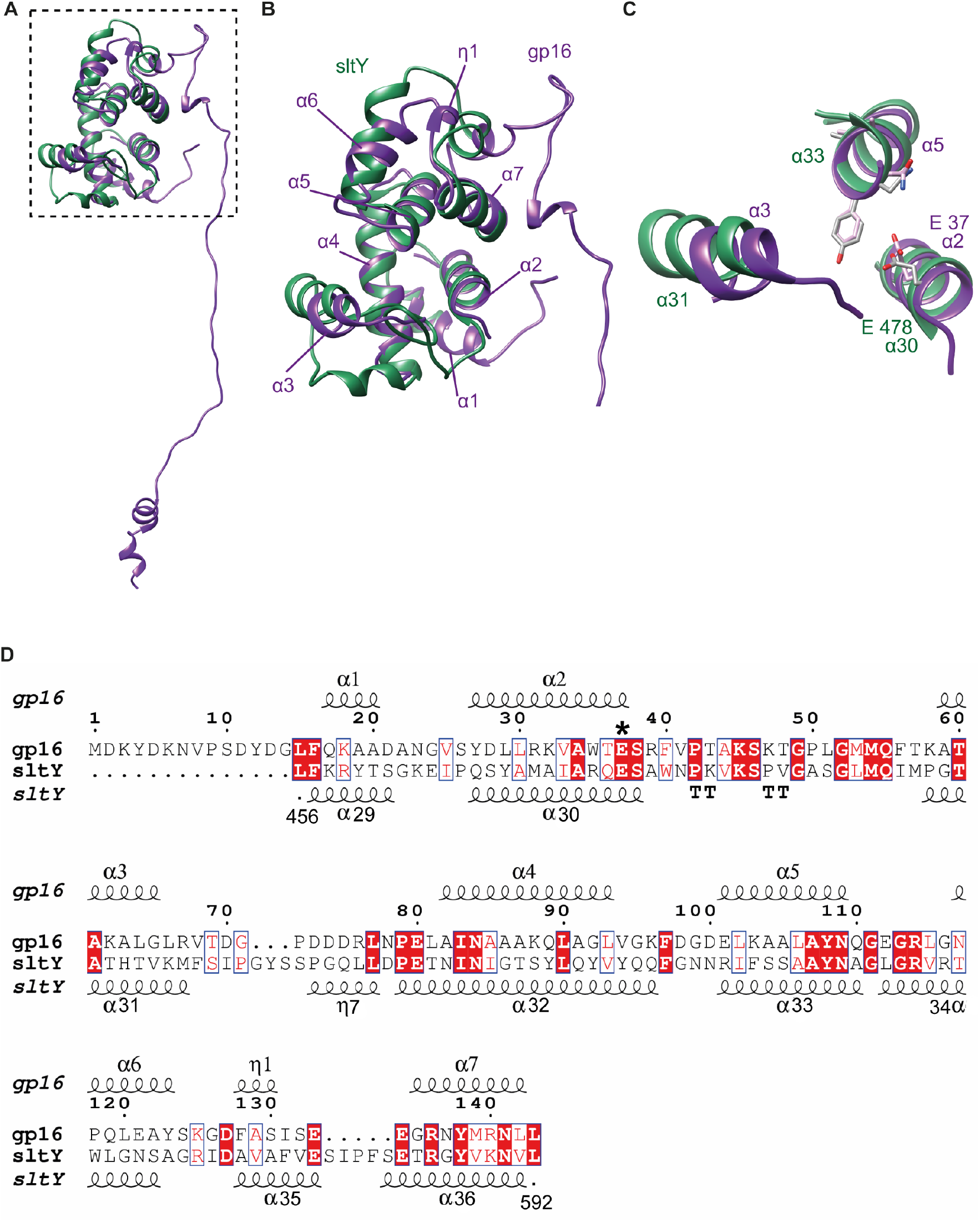
The T7 gp16 transglycosylase domain. **A.** Ribbon representation of the reconstructed part of the gp16 monomer when bound to gp15 (purple), docked into the *E. coli* SltY lytic transglycosylase domain (PDB 1SLY, green). **B.** Zoom-in of the dashed square showed in A, with arrows pointing to the gp16 α-helices. **C.** Overlapping of gp16 and SltY helices involved in the active site: gp16 helices α2, α3 and α5 (purple) and SltY helices α30, α31 and α33 (green). Conserved gp16 residues (in grey): E37 (catalytic glutamate), A107, Y108, N109; and conserved SltY residues (in pink): E478 (catalytic glutamate), A551, Y552, N553, A554, are shown in a stick representation. **D.** Sequence alignment of gp16 (from M1 to L146) and *E. coli* SltY genes (from L456 to L592) obtained with ESPrit 3 software (47); homologous residues are highlighted in red, both catalytic glutamate are shown with an asterisk. Secondary structure elements and residue number for both proteins are also indicated.

As described, the largest visible part of gp16 in the gp15-gp16 complex is the transglycosylase domain. This domain consists of 7 α-helices and as expected, docking of this domain into the *E. coli* SltY catalytic domain showed a good overlapping of the two structures (Fig. 3A-C) (17, 23), from L15 in helix α1 to L143 in helix α7 (RMSD 0.951 Å). There are some regions whose overlap is slightly worse (Fig. 3B): the gp16 helices α3 and helix α5 are inclined around 10° with respect to SltY helices α31 and α33; as it was mentioned before the loops between gp16 helix α2 to helix α3 (residues F40-M54) and helix α3 to helix α4 (residues L66-R76) were not properly defined and could not be traced; and the loop between gp16 helix α5 and helix α6 is shorter in the case of gp16 and lies in a different position.

The analysis of both proteins demonstrated that gp16 and SltY share the same structural arrangement of the active site, including the well characterized catalytic residue E37 found at the end of helix α2 (16, 17). E37 in α2 overlaps with the active-site amino acid SltY-E478 in α30 (Fig 3C), confirming the main role of this residue in peptidoglycan degradation during the infection process (16). Concerning the conserved residues important for the architecture of the active site in SltY (17, 24), while the GXMQ site is not well observed in gp16, the AYNXG site (in gp16 helix α5) shows a perfect overlapping (Fig. 3B and C). Surprisingly, the residues in SltY helix α33 involved in the interaction with the peptidoglycan (17, 24) do not show either exactly the same position than their gp16 counterparts. As it was mentioned before, its equivalent helix (gp16 helix α5) is turned around 10° with respect to the one of StlY (Fig. 3B), which suggests some structural flexibility in this region. The existence of certain flexibility in the transglycosilase domain is also supported by the lower resolution of this area compared to the rest of the structure (Supplementary Fig. 4E). We suggest that a certain flexibility might be needed in this domain to perform its biological function as a peptidoglycan-degrading enzyme.

### Structural characterization of the interactions that stabilize the gp15-g16 core complex

We then analyzed in detail the interactions that induce stabilization of gp15 upon interaction with a small part of gp16 (Fig. 4). This interaction is of a staggered nature, with one subunit of gp16 interacting with two subunits gp15 subunits: on the one hand the transglycosilase domain and the loop domain of a gp16 molecule interact with the HBD2 and the α15 domains of one gp15 subunit (Fig. 4 A, B and C) and on the other hand the continuation of the loop and the tip domain of the same gp16 molecule interact with the HBD3 and the base domain of the adjacent gp15 subunit (Fig. 4D-F). Gp15 builds the inner core of the complex and presents the highest inter-monomeric surface of interaction with 36,715 A^2^ of inter-subunit contacts. In contrast, the gp16 subunits hardly interact with each other in the resolved structure (Supplementary Fig. 4E), and they depend on the formation of an initial gp15 hexamer to assemble into the complex. This could explain why gp16 is found in solution as a monomer, regardless of the concentration tested (Supplementary Fig. 1D). Gp16 interacts with gp15 through the transglycosylase and tip domains, but the most remarkable interaction is though its 50-residue loop, which follows closely the new folded domains of gp15: the end of the α-helix 15 (Fig. 4B and C) as well as the HBD3 the base domains of the adjacent gp15 subunit (Fig 4 E and F). These interactions, based on hydrophobic contacts (Fig. 4 B and E), and in salt bridges and hydrogen bonds (Fig 4 C and F), are probably key in assisting the complete folding of gp15, and thus in the gp15-gp16 complex formation.

**Figure 4.**
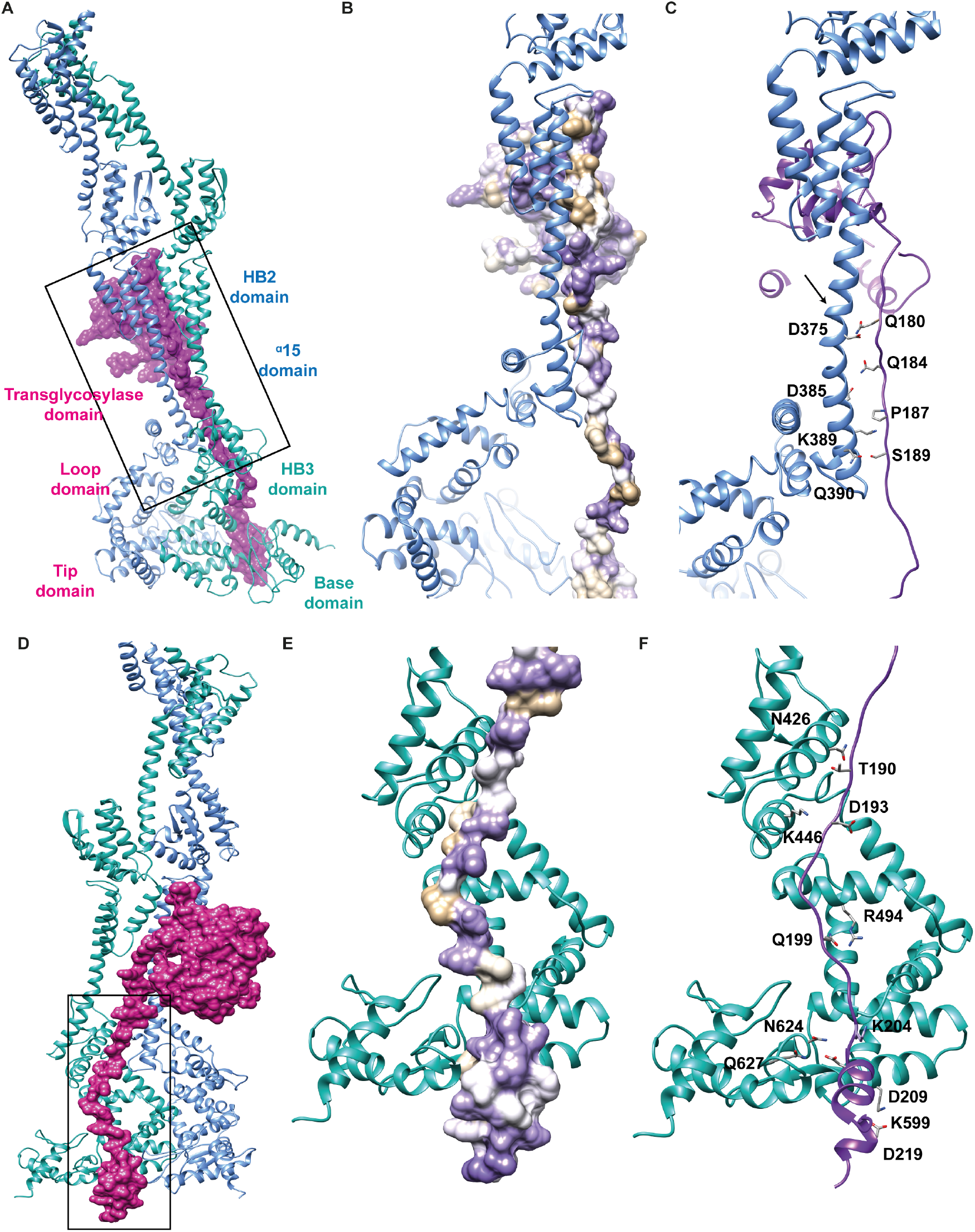
Structural analysis of the gp15-gp16 complex interaction. **A, D.** Representation of two subunits of gp15 (in ribbon blue and turquoise) and one subunit of gp16 (in surface purple) showing the interaction of one gp16 subunit with two gp15 molecules in two different orientations. In A the location of the domains is labeled. **B, E. A** detailed view of the square depicted in A and D, respectively showing the gp16 hydrophobic cluster involved in the interaction between gp15 α15 (B) and base domain (E) (ribbon) and gp16 loop and tip domain (hydrophobic surface representation). Gp16 hydrophobic surface was calculated with Chimera software (47) according to the Kyte and Doolittle scale (48), tan represents more hydrophobic residues and purple polar residues. **C, F.** The same view in B and E showing one subunit of gp15 (blue and turquoise) and gp16 (purple) in ribbon representation. In **C**, the residues involved in hydrogen bond and salt bridge formation between gp15 α15 (D375, D385, K389, Q390) and gp16 loop (Q180, Q184, P187, S189) are indicated. The position of the last residue of α15 gp15 folded in the gp15 structure solved alone is indicated with an arrow. In **F,** the residues involved in hydrogen bond and salt bridge formation between gp15 HBD3 and base domains (N426, K446, R494, K599, N624, Q627) and gp16 loop and tip (T190, D193, Q199, K204, D209, D219) are indicated.

## DISCUSSION

The mechanism of DNA ejection of bacteriophage T7 has been the subject of intense study and controversy for many years. The T7 core proteins assemble inside the capsid as a toroidal structure surrounded by the DNA (13), and it has been postulated that they unfold and refold as a tubular structure in the periplasmic space (11). In this article, we describe for the first time the structure of the gp15-gp16 complex, formed by the two main components of the gp14-gp15-gp16, core complex. The atomic model of gp15-gp16 solved by cryo-EM shows that these two T7 core proteins are able to assemble into a tubular structure, supporting the hypothesis that T7 can build an ‘extensible tail’. The length of the core complex (around 210 Å) is compatible with the dimensions of the bacterial periplasmic space. In order to check this hypothesis, we docked our 3D reconstruction of core complex gp15-gp16 *in vitro* into the cryoelectron tomography reconstruction of bacteriophage T7 obtained by Hu and colleagues (11) (Fig. 5A). Docking of our gp15-gp16 complex into the corresponding part of the bacteriophage reconstruction is compatible with the overall dimensions of the previously solved structure (11), suggesting that our 3D model assembled *in vitro* could be the T7 translocation conduit for DNA ejection. The structure of the gp15-gp16 complex has a channel with an overall diameter of 30 Å, which can accommodate a dsDNA molecule. Nevertheless, the entrance and exit of the channel are narrowed (21–24 Å, Fig. 2C), thus suggesting that the gp15-gp16 core complex characterized in this work could be an intermediate translocation complex.

**Figure 5.**
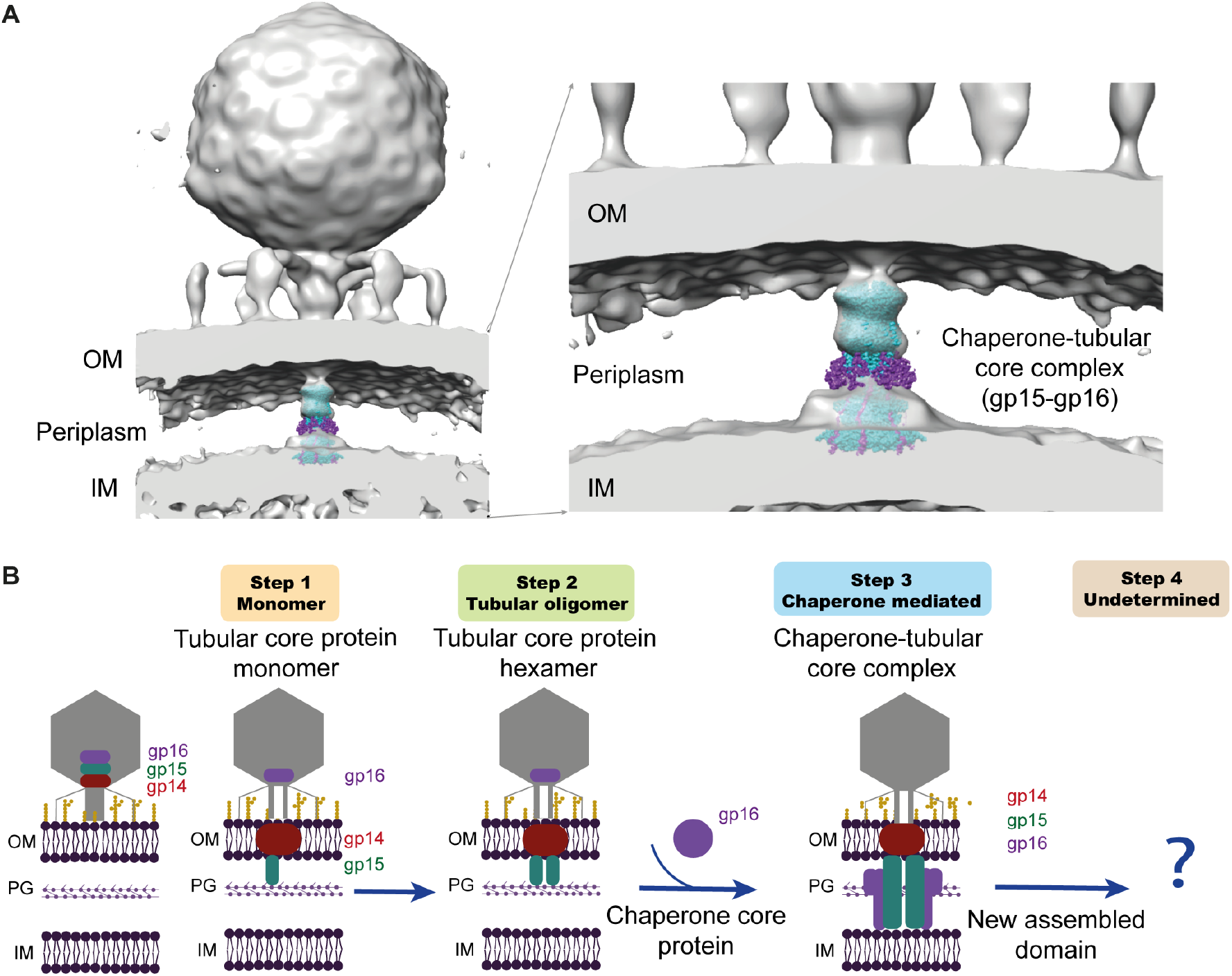
Assembly of the T7 translocation complex in the bacterial periplasmic space. **A.** Docking of the 3D reconstruction of the core complex assembled *in vitro* (gp15 in light blue, gp16 in purple) into the 3D model of bacteriophage T7 during DNA ejection obtained by cryo-ET in Hu et al. ^11^. **B.** Schematic of the proposed assembly model of the gp15 and gp16 core complex during infection, showing T7 bacteriophage (grey); *E. coli* bacterial wall composed by the internal membrane (IM), outer membrane (OM), and peptidoglycan (PG) and the assembly of the viral proteins gp14 (red), gp15 (green) and gp16 (purple). The three proteins forms part of the core complex, a structured assembly present at the vertex of the mature phage. After interaction with the bacterial membrane, the three proteins undergo a sequential unfolding to traverse the phage tail. Gp14 refolds and forms an oligomer that allows the passing of gp15 through the outer membrane (step 1). Gp15 hexamerizes part of its structure (step 2) and needs the presence of gp16 to form the whole hexagonal, tubular structure with long enough to reach the bacterial inner membrane (step 3). Although most of gp16 is still unstructured, a transglycosilase domain has been formed that hydrolyzes the PG layer. A yet unknown factor is needed for the total assembly of gp16 and thus the formation fully structured and active periplasmic core complex ready for the translocation of the viral genome (step 4).

Although only 206 residues of the gp16 protein were solved in our translocation complex, this structure shows the key interactions that are required for the full folding of gp15, and also the presence of a canonical transglycosylase domain (Fig 3), which shares the same structural arrangement of the active site with sltY *E. coli* transglycosylase enzyme (Fig. 3C). Observation of the full gp15-gp16 complex size by AUC strongly suggests that the gp16 missing region in this work is disordered and thus not observed in our structural studies. It is important to reinforce the notion that that due to their biological function, gp15 and g16 proteins, which are fully structured in the mature virus and form part of the core complex (13), are able to unfold to transverse the viral tail channel and reach the periplasm, and we have captured in part this state for gp15 (which form hexamers on its own in which only half of the sequence is structured) and gp16 (which is non-structured on its own). The interaction of gp15 and gp16 in the periplasm induces the folding of the whole gp15, and further interactions with other phage proteins (such as gp14) or bacterial membrane components may be essential to promote the fully assembly of gp16,, and to trigger the conformational changes needed in the complex to ensure a fully open translocation path.

The intermediate structures characterized in this work allow us to propose a model for the assembly of gp15 and gp16 proteins outside the phage structure during T7 infection (Fig. 5B). This two proteins, together with gp14, form the core complex of the mature virus (Fig. 5B). The interaction of the tail fibers with the T7 LPS receptor triggers the conformational changes needed for the tail opening and subsequent capsid core disassembly (19). Previous work suggests that gp14 is the first one to exit the viral particle to build a pore in the outer bacterial membrane (OM) (Fig. 5B, step 1) (18, 25). Monomers of gp15 are then released from the viral capsid into the *E. coli* periplasmic space and interact with the gp14 pore, probably through the N-terminal, electropositive area of gp15 (Fig. 2G). In a following step, the gp15 monomers oligomerize in a partially folded hexameric tube (Fig. 5B, step 2). Subsequently, the arrival of gp16 and its assembly to the gp15 hexamer generates the gp15-gp16 complex and the transglycosylase motifs for bacterial peptidoglycan degradation, through the puncturing of the peptidoglycan layer (Fig. 5B, step 3). Gp16 acts as a chaperone by assisting the folding of the full structure of the gp15 hexamer that now extends its tubular structure to reach the inner bacterial membrane (IM). In a final, not yet structurally characterized step (Fig. 5B, step 4), the partially folded gp15-gp16 complex could interact with the inner bacterial membrane (IM), specially, thanks to the gp16 property to associate with lipid bilayers (18). This it could allow the formation the fully structured and active periplasmic core complex ready for the translocation of the viral genome. The lack of structure in the preliminary steps of both gp15 and gp16 could permit their exit from the phage head and to traverse the gp14 pore, and only after a proper interaction (gp15 with gp16, gp16 with a yet unknown molecule), reach their final, structured conformation.

## METHODS

### Protein expression and purification

The bacteriophage T7 *gp15* gene was inserted into the pRSET-B vector (amp^r^ and 6x-His tag) from Invitrogen (USA), between the *Xho*I and *EcoR*I restriction sites. The gp15 protein was overexpressed in *Escherichia coli* C41 (DE3) grown in LB media supplemented with 100 mg/ml ampicillin at 37 °C, with induction by 1 mM isopropyl-b-D-1-thiogalactopyranoside (IPTG) for 3h when an optical density (O.D._600_) of 0.4-0.6 was reached. Bacterial cells were resuspended in 100 mM sodium citrate dihydrate, 10 mM MgCl_2_, 1 mM phenylmethane sulfonyl fluoride (PMSF) 100 mM Tris pH 8.0, and Complete Protease Inhibitor Cocktail Tablets (Roche) and lysed using a cell disruptor. Cell debris was discarded after centrifugation for 15 min at 3,000 x g. After that, gp15 was purified following two chromatographic steps. Briefly, the protein was purified, always at 10 °C and, loaded onto a HisTrap HP column equilibrated in buffer A: 100 mM Tris pH 8.0, 100 mM sodium citrate dihydrate, 10 mM MgCl_2_, 1 mM phenylmethane sulfonyl fluoride (PMSF) and eluted in buffer A with 250 mM imidazole. The second step of purification consisted of a size exclusion chromatography (SEC) using a Superdex200 HiLoad column equilibrated in buffer A. Finally, the purified protein was concentrated using Amicon Ultra 50 k (Millipore).

The bacteriophage T7 *gp16* gene was cloned into the pET-16b vector (amp^r^ and 10x-His tag) from Novagen (Germany), inserted between *Nde*I and *Xho*I restriction sites. Overexpression of gp16 in *E. coli* C41 (DE3) was carried out under specific conditions due to the toxic effects of its transglycosylase activity. The culture was grown at 37 °C using autoinduction-media (26). Afterwards, when the O.D._600_ reached ~0.5, the culture was transferred to 16 °C for 48 h. The protein was harvested and purified following the same steps described for gp15 above.

The gp15-gp16 core complex was obtained *in vitro* after incubation of the purified proteins for 30 min at room temperature (RT) in a thermo block with shaking (Eppendorf). The proteins were incubated using 180 pmol each at a 1:1 ratio. The complexes were purified in a 10-40% (v/v) glycerol gradient in buffer A, the gradient was formed into Gradient Station Biocomp station with 53s/86°/20 rpm. The gradient was stored ~1 h at 4 °C and then the complexes were separated by ultracentrifugation at 165,500 xg for 17 h using a SW55 Ti rotor at 10 °C. The fractions were collected from bottom to top and checked in a 10% SDS-Page gel.

### Sample checking by negative staining electron microscopy

After obtaining pure and suitable gp15, gp16 and gp15-gp16 core proteins, a preliminary electron microscopy evaluation was performed using negative staining. The samples were transferred to an 80kV JEM1010 (Jeol) with a CMOS 4×4 sensor (TemCam F416) yielding a pixel size of 2.5 Å/pix at the Electron Microscopy Services of the Centro de Biología Molecular Severo Ochoa (CBMSO-CSIC), and data processing was performed using the same workflow in each sample, using the Scipion software framework (27). The contrast transfer function (CTF) was estimated using ctffind4 (28). Particles were manually picked using XMIPP (29) and 2D and 3D classification was performed using RELION, the latter applying C6 symmetry (30). A total of 1648 and 2553 particles were used to 3D reconstruct gp15 and the gp15-gp16, respectively, at a low resolution (~ 20-22 Å).

### Cryo-EM sample preparation and data acquisition

Grids containing either gp15 or the gp15-gp16 core complex were vitrified using a Thermo Fisher Vitrobot Mark IV. R2/2 Quantifoil grids were glow-discharged for 1 min and loaded with 3 μl of purified sample at around 1 mg/ml for gp15 and 1.5 mg/ml for the core complex. Samples of gp15 and the gp15-gp16 complex were then incubated for 3 or 1 min, respectively, at 95% humidity and 22 °C inside the Vitrobot, cleaned and blotted twice in buffer A to remove the excess glycerol in the buffer prior to freezing. Grids were then blotted for 3.5 s with forces between −5 and −7 and plunge-frozen in liquid ethane.

The gp15 grids was transferred to a Talos Arctica (Thermo Fisher) electron microscope operated at 200 kV (at the cryo-EM Service, CNB, Spain), and 2,470 movies fractioned in 62 frames were recorded in an automated fashion on a Falcon III (Thermo Fisher) detector using EPU (Thermo Fisher) with a pixel size of 0.855 Å/pix; fraction exposure time was ~35 s and total accumulated dose ~30 e^−^/Å^2^ (~0.48 e^−^/Å^2^ /frame) (Supplementary Table 1).

For the gp15-gp16 complex, the grids were transferred to a Titan Krios (Thermo Fisher) electron microscope operated at 300 kV (at the eBIC, Diamond Light Source, UK). A total of 5,506 movies fractioned in 32 frames were collected on a K2 Summit detector (Gatan) using EPU (Thermo Fisher) with a pixel size of 1,047 Å/pix, 8 s of fraction exposure time and total accumulated dose ~40 e^−^/Å^2^ (~1.25 e^−^/Å^2^ /frame) (Supplementary Table 1).

### Image processing and map calculation

Cryo-EM data processing was performed using the Scipion software framework (27). Dose-fractionated image stacks were motion-corrected and dose-weighted using MotionCor2 (31). Contrast transfer function (CTF) was estimated using ctffind4 and xmipp3 software (28, 29) for the gp15 data set and GCTF software (32) for the core complex data set. Particles of gp15 were picked using Gautomatch (developed by Zhang K, MRC-LMB Cambridge, UK) and the gp15-gp16 core complex using RELION (30). The particles extracted were classified using RELION 2D and 3D (30, 33) and the initial volume was built using Ransac (29). C6 symmetry was applied for gp15 and the gp15-gp16 complex. The volumes were obtained using RELION 3D auto-refine (34) with 50,980 and 72,882 particles for gp15 and the gp15-gp16 complex, respectively. In the case of the gp15-gp16 data, multiple rounds of CTF refinement and Bayesian polishing (35) were performed before the final 3D autorefinement and postprocessing with RELION. In all the cases, structure resolutions were estimated from RELION FSC curves applying the 0.143 cutoff criteria (30, 36, 37), and local resolutions were computed with MonoRes (38). Final volumes for gp15 and the gp15-gp16 complex were respectively post-processed for local sharpening with LocaldeBlur (39), using MonoRes (38) volume as input.

### Model building and coordinate refinement

Long-version of gp15 tubular core protein present in the gp15-gp16 complex was traced *ab initio* with Coot (40), using the PSIPRED server (41) to predict secondary structure as a guide. This structure was then used as a template to trace the short version of the partially folded gp15. Gp16 was firstly traced *ab initio* with Coot using the PSIPRED (41) secondary structure prediction information, this initial model was used as input in DALI server (21), allowing to identify the homology of the traced region of gp16 with SltY (1sly) and Mltc (4cfo) transglycosylases that were then used as a guide to built the final gp16 model. In both cases, atomic models were refined using PHENIX real space refinement (42). Refinement and validation scores are shown in Supplementary Table 1.

### Sedimentation velocity assays (SV)

Sedimentation velocity assays (SV) were performed at the Molecular Interaction Facility at the Centro de Investigaciones Biológicas Margarita Salas (CIB-CSIC). 320 μl aliquots of gp15, gp16 and the gp15-gp16 complex were loaded onto analytical ultracentrifugation cells at concentrations of 1, 0.5 and 0.2 mg/ml in 100 mM sodium citrate dihydrate, 10 mM MgCl_2_ 100 mM Tris, pH 8.0 buffer. The experiments were carried out at 10 °C and 43,000 rpm in a XL-I analytical ultracentrifuge (Beckman-Coulter Inc.) equipped with both UV-VIS absorbance and Raleigh interference detection systems, using an An-50Ti rotor, and 12 mm Epon-charcoal standard double-sector centerpieces. Sedimentation profiles were recorded at 280 nm, and differential sedimentation coefficient distributions were calculated by least-squares boundary modeling of sedimentation velocity data using the continuous distribution c(*s*) Lamm equation model as implemented by SEDFIT (43). These *s* values were corrected to standard conditions (water, 20 °C, and infinite dilution) (44) using the program SEDNTERP (45) to get the corresponding standard *s* values (*s*_20,*w*_).

### Dynamic light scattering assays (DLS)

DLS experiments were also carried out at the Molecular Interaction Facility at the Centro de Investigaciones Biológicas Margarita Salas, (CIB-CSIC). The experiments were performed using a Protein Solutions DynaPro MS/X instrument (Protein Solutions, Piscataway, NJ) at 20 °C and a 90° light scattering cuvette. The samples, under the same experimental conditions used in AUC, were centrifuged for 10 min at 12,000 xg and 21 °C just before measurements. Data were collected and analyzed with Dynamics V6 Software.

### Estimation of molar mass of the core proteins from hydrodynamic measurements

The apparent molar mass of single sedimenting solute species (*M*) were calculated using measured values of the sedimentation coefficient *s* and the diffusion coefficient *D* according to the Svedberg equation (46). The estimated molar mass obtained using this relation is independent of the shape of the sedimenting/diffusing species, as frictional coefficients for sedimentation and diffusion cancel in the derivation of the equation.

## Data availability

The electron microscopy maps were deposited in the Electron Microscopy Data Bank (EMDB) under accession codes EMD-10911 and EMD-10912. The atomic coordinates structure factors were deposited in the Protein Data Bank (PDB) under accession codes 6YSZ and 6YT5. All relevant data are available from the authors upon request.

## Acknowledgements

This work was supported by the Ministry of Science, Innovation and Universities of Spain, grant BFU 2014-54181 to JLC, contracts SEV-2013-0347 to AC and BES-2015-073615 to MPR. The authors acknowledge the support and use of resources of Instruct-ERIC for image processing. The authors wish to thank the personnel of the following facilities for help during electron microscopy checking, cryo-EM and molecular interactions data acquisition: the Electron Microscopy Services of the Centro de Biología Molecular Severo Ochoa (CBMSO-CSIC) and the Centro Nacional de Biotecnología (CNB-CSIC) in Madrid (Spain); the Cryo-EM Facility of the Centro Nacional de Biotecnología-Centro de Investigaciones Biológicas Margarita Salas (CNB-CIB) in Madrid (Spain); the Molecular Interactions Facility of the Centro de Investigaciones Biológicas (CIB-CSIC) in Madrid (Spain) and the Electron Bio-Imaging Centre (eBIC) at Diamond Light Source in Didcot (UK). The authors are grateful to Erney Ramírez-Aportela for his help in the application of sharpening methods to improve the density of the gp15-gp16 complex map. The professional editing service NB Revisions was used for technical preparation of the text prior to submission.

## Author contributions

JLC and AC designed and coordinated the project. AC, MPC and MPR prepared the proteins. JRLO collected and processed the molecular interaction data. AC and MPR collected, processed the cryo-EM data and solved the cryo-EM structures. JLC, AC, JMV and MPR wrote the paper.

## References

1. A. Bertin, M. de Frutos, L. Letellier, Bacteriophage-host interactions leading to genome internalization. Curr Opin Microbiol 14, 492–496 (2011).

2. A. A. Aksyuk, M. G. Rossmann, Bacteriophage assembly. Viruses 3, 172–203 (2011).

3. F. L. Nobrega et al., Targeting mechanisms of tailed bacteriophages. Nat Rev Microbiol 16, 760–773 (2018).

4. A. Cuervo, J. L. Carrascosa, Viral connectors for DNA encapsulation. Curr Opin Biotechnol 23, 529–536 (2012).

5. S. R. Casjens, The DNA-packaging nanomotor of tailed bacteriophages. Nat Rev Microbiol 9, 647–657 (2011).

6. S. R. Casjens, I. J. Molineux, Short noncontractile tail machines: adsorption and DNA delivery by podoviruses. Adv Exp Med Biol 726, 143–179 (2012).

7. L. Sun et al., Icosahedral bacteriophage PhiX174 forms a tail for DNA transport during infection. Nature 505, 432–435 (2014).

8. B. Peralta et al., Mechanism of membranous tunnelling nanotube formation in viral genome delivery. PLoS Biol 11, e1001667 (2013).

9. A. Bhardwaj, A. S. Olia, G. Cingolani, Architecture of viral genome-delivery molecular machines. Curr Opin Struct Biol 25, 1–8 (2014).

10. C. Wang, J. Tu, J. Liu, I. J. Molineux, Structural dynamics of bacteriophage P22 infection initiation revealed by cryo-electron tomography. Nat Microbiol 10.1038/s41564-019-0403-z (2019).

11. B. Hu, W. Margolin, I. J. Molineux, J. Liu, The bacteriophage t7 virion undergoes extensive structural remodeling during infection. Science 339, 576–579 (2013).

12. A. Cuervo et al., Structures of T7 bacteriophage portal and tail suggest a viral DNA retention and ejection mechanism. Nat Commun 10, 3746 (2019).

13. X. Agirrezabala et al., Maturation of phage T7 involves structural modification of both shell and inner core components. EMBO J 24, 3820–3829 (2005).

14. G. S. Roeder, P. D. Sadowski, Bacteriophage T7 morphogenesis: phage-related particles in cells infected with wild-type and mutant T7 phage. Virology 76, 263–285 (1977).

15. M. E. Cerritelli, J. F. Conway, N. Cheng, B. L. Trus, A. C. Steven, Molecular mechanisms in bacteriophage T7 procapsid assembly, maturation, and DNA containment. Adv Protein Chem 64, 301–323 (2003).

16. M. Moak, I. J. Molineux, Peptidoglycan hydrolytic activities associated with bacteriophage virions. Mol Microbiol 51, 1169–1183 (2004).

17. M. Moak, I. J. Molineux, Role of the Gp16 lytic transglycosylase motif in bacteriophage T7 virions at the initiation of infection. Mol Microbiol 37, 345–355 (2000).

18. S. Leptihn, J. Gottschalk, A. Kuhn, T7 ejectosome assembly: A story unfolds. Bacteriophage 6, e1128513 (2016).

19. V. A. Gonzalez-Garcia et al., Conformational changes leading to T7 DNA delivery upon interaction with the bacterial receptor. J Biol Chem 290, 10038–10044 (2015).

20. A. J. Noble et al., Reducing effects of particle adsorption to the air-water interface in cryo-EM. Nat Methods 15, 793–795 (2018).

21. L. Holm, DALI and the persistence of protein shape. Protein Sci 29, 128–140 (2020).

22. H. Engel, B. Kazemier, W. Keck, Murein-metabolizing enzymes from Escherichia coli: sequence analysis and controlled overexpression of the slt gene, which encodes the soluble lytic transglycosylase. J Bacteriol 173, 6773–6782 (1991).

23. A. M. Thunnissen et al., Doughnut-shaped structure of a bacterial muramidase revealed by X-ray crystallography. Nature 367, 750–753 (1994).

24. E. J. van Asselt, A. M. Thunnissen, B. W. Dijkstra, High resolution crystal structures of the Escherichia coli lytic transglycosylase Slt70 and its complex with a peptidoglycan fragment. J Mol Biol 291, 877–898 (1999).

25. C. Y. Chang, P. Kemp, I. J. Molineux, Gp15 and gp16 cooperate in translocating bacteriophage T7 DNA into the infected cell. Virology 398, 176–186 (2010).

26. F. W. Studier, Protein production by auto-induction in high density shaking cultures. Protein Expr Purif 41, 207–234 (2005).

27. J. M. de la Rosa-Trevin et al., Scipion: A software framework toward integration, reproducibility and validation in 3D electron microscopy. J Struct Biol 195, 93–99 (2016).

28. A. Rohou, N. Grigorieff, CTFFIND4: Fast and accurate defocus estimation from electron micrographs. J Struct Biol 192, 216–221 (2015).

29. J. M. de la Rosa-Trevin et al., Xmipp 3.0: an improved software suite for image processing in electron microscopy. J Struct Biol 184, 321–328 (2013).

30. S. H. Scheres, RELION: implementation of a Bayesian approach to cryo-EM structure determination. J Struct Biol 180, 519–530 (2012).

31. S. Q. Zheng et al., MotionCor2: anisotropic correction of beam-induced motion for improved cryo-electron microscopy. Nat Methods 14, 331–332 (2017).

32. K. Zhang, Gctf: Real-time CTF determination and correction. J Struct Biol 193, 1–12 (2016).

33. S. H. Scheres, Processing of Structurally Heterogeneous Cryo-EM Data in RELION. Methods Enzymol 579, 125–157 (2016).

34. J. Zivanov et al., New tools for automated high-resolution cryo-EM structure determination in RELION-3. Elife 7 (2018).

35. J. Zivanov, T. Nakane, S. H. W. Scheres, A Bayesian approach to beam-induced motion correction in cryo-EM single-particle analysis. IUCrJ 6, 5–17 (2019).

36. S. H. Scheres, S. Chen, Prevention of overfitting in cryo-EM structure determination. Nat Methods 9, 853–854 (2012).

37. P. B. Rosenthal, R. Henderson, Optimal determination of particle orientation, absolute hand, and contrast loss in single-particle electron cryomicroscopy. J Mol Biol 333, 721–745 (2003).

38. J. L. Vilas et al., MonoRes: Automatic and Accurate Estimation of Local Resolution for Electron Microscopy Maps. Structure 26, 337–344 e334 (2018).

39. E. Ramirez-Aportela et al., Automatic local resolution-based sharpening of cryo-EM maps. Bioinformatics 10.1093/bioinformatics/btz671 (2019).

40. P. Emsley, K. Cowtan, Coot: model-building tools for molecular graphics. Acta Crystallogr D Biol Crystallogr 60, 2126–2132 (2004).

41. D. W. Buchan, F. Minneci, T. C. Nugent, K. Bryson, D. T. Jones, Scalable web services for the PSIPRED Protein Analysis Workbench. Nucleic Acids Res 41, W349–357 (2013).

42. P. V. Afonine et al., Real-space refinement in PHENIX for cryo-EM and crystallography. Acta Crystallogr D Struct Biol 74, 531–544 (2018).

43. P. Schuck, Size-distribution analysis of macromolecules by sedimentation velocity ultracentrifugation and lamm equation modeling. Biophys J 78, 1606–1619 (2000).

44. K. E. Holde, W. C. Johnson, P. S. Ho, in Principles of Physical Biochemistry. (Pearson Prentice Hall., 1985).

45. T. Laue, B. Shah, T. Ridgeway, S. Pelletier, “Computer-aided interpretation of analytical sedimentation data for proteins” in Analytical Ultracentrifugation in Biochemistry and Polymer Science, S. Harding, A. Rowe, J. Horton, Eds. (Royal Society of Chemistry; Cambridge, 1992), pp. 90–125.

46. T. Svedberg, K. O. Pedersen, the ultracentrifuge. (Claredon Press, Oxford., 1940).

47. X. Robert, P. Gouet, Deciphering key features in protein structures with the new ENDscript server. Nucleic Acids Res 42, W320–324 (2014).

48. J. Kyte, R. F. Doolittle, A simple method for displaying the hydropathic character of a protein. J Mol Biol 157, 105–132 (1982).

