## Supplementary Figures and Table for "Assisted assembly of bacteriophage T7 core components for genome translocation across the bacterial envelope"

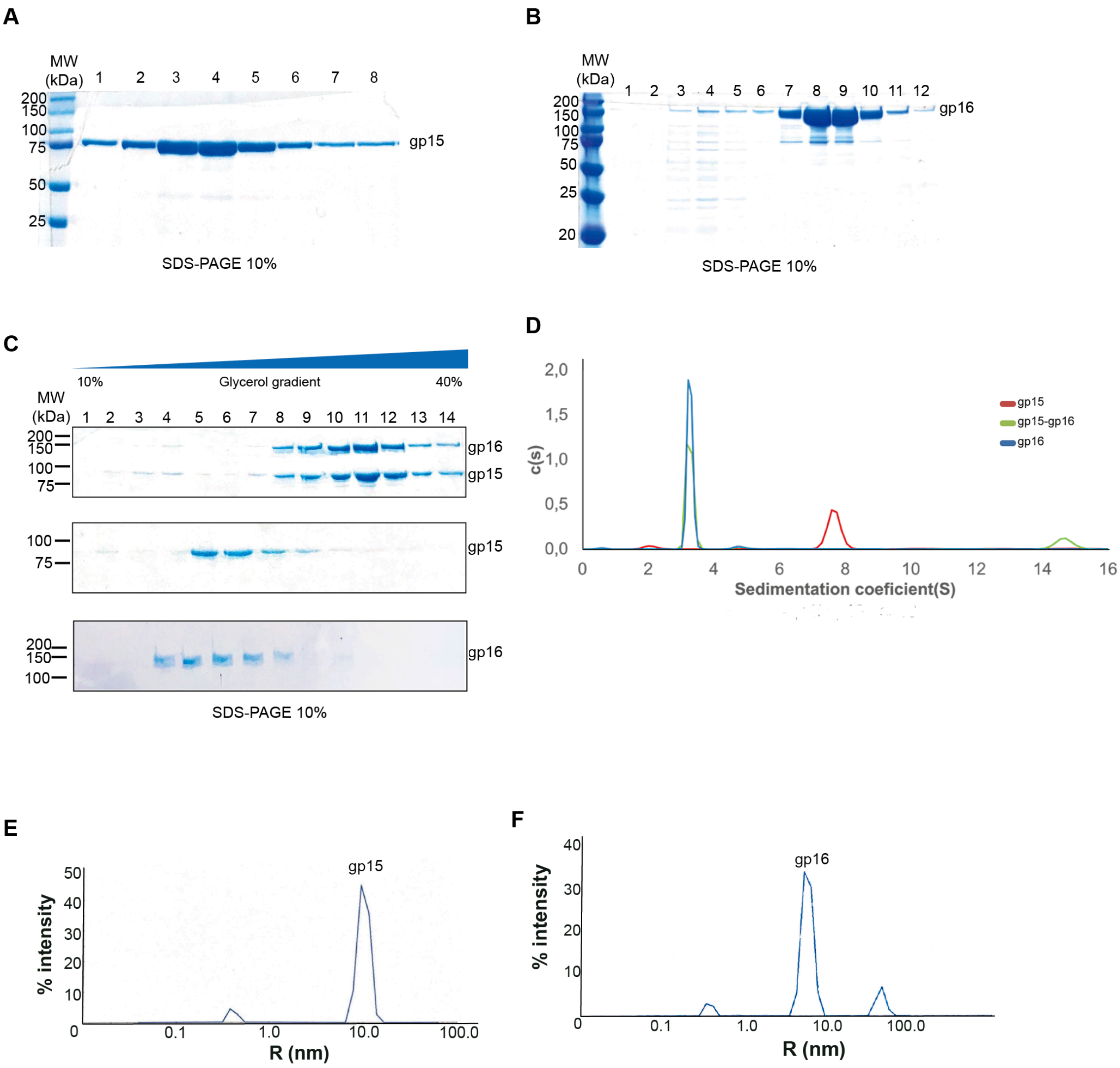

**Supplementary Figure 1. The protein purification process for gp15, gp16 and gp15-gp16 complex, and the analytical ultracentrifugation (AUC) by sedimentation velocity (SV) of the core proteins. A.** 10% SDS-PAGE gel of gp15 size exclusion chromatography purification fractions, stained with Bio-Safe Coomassie. Lanes: MW, markers of molecular weight (Precision Plus Protein TM dual color standards); from 1 to 8, eluted gp15 protein fractions. **B.** 10% SDS-PAGE gel showing size exclusion fractions stained with Bio-Safe Coomassie; fractions from 7-12 correspond to the eluted gp16 protein. **C**. 10% SDS-PAGE gel of gp15-gp16 complex fractions (1 to 14) stained with Bio-Safe Coomassie. The direction of the 10-40% (v/v) glycerol gradient and the position of gp15 and gp16 proteins is indicated. **D.** Overlapping of all the core protein SV profiles from isolated proteins gp15 (red), gp16 (blue) and gp15-gp16 complex (green). The rotor speed was 43,000 rpm at 10º C and sedimentation profiles were recorded at 280 nm. **E.** Dynamic light scattering (DLS) assay of gp15 protein. The hydrodynamic radius distribution by intensity for gp15 at 1 mg/ml. DLS plot showing hydrodynamic radius distribution (R) and the intensity. The major peak corresponds to R=10.3 nm and the average of translational diffusion coefficient D is 1.51 x $10^{-11}$ $m^2/s$. **F**. DLS assay of gp16 protein at 1 mg/ml.

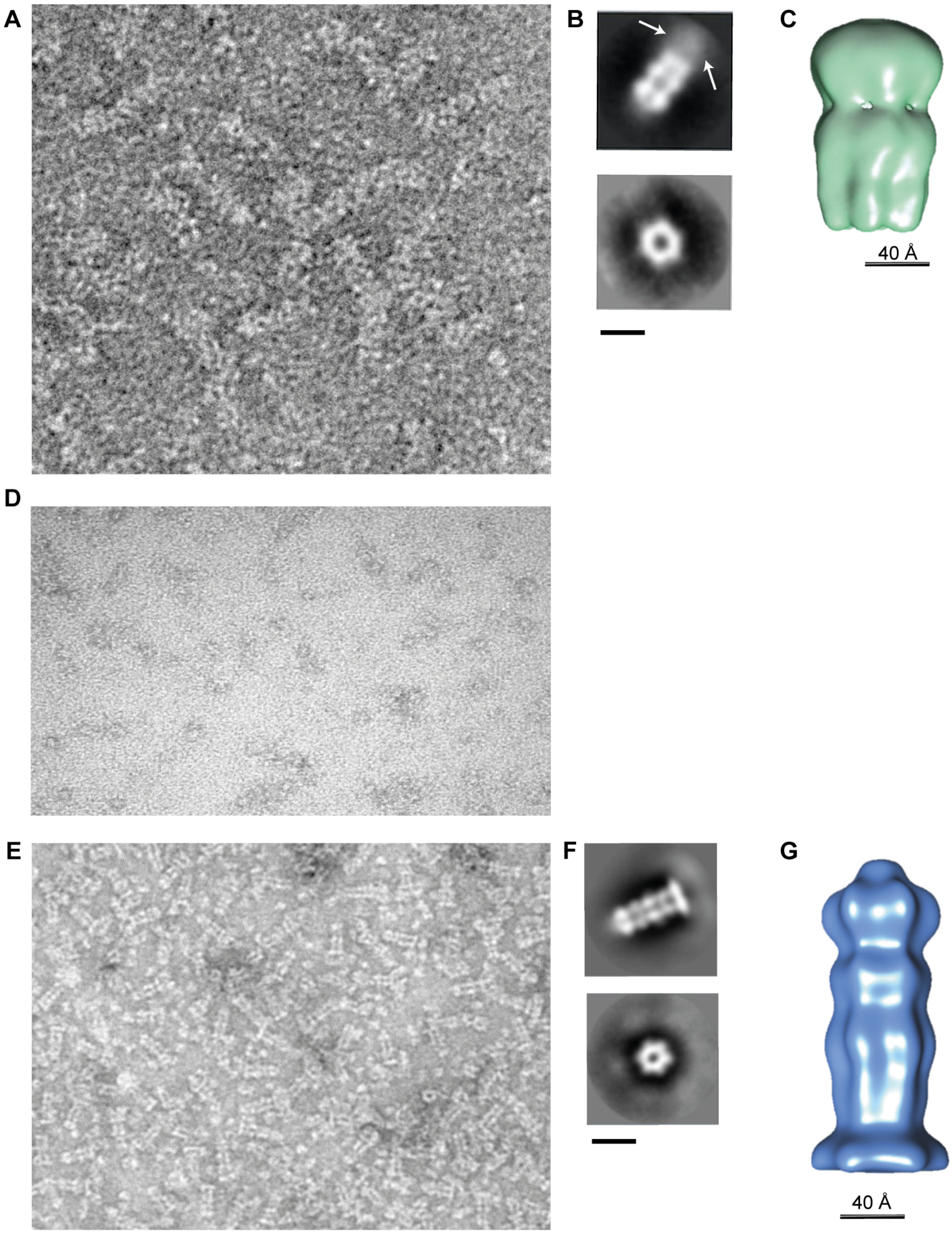

**Supplementary Figure 2. Negative staining of the gp15, gp16 and gp15-gp16 core proteins. A.** A representative gp15 micrograph obtained using an 80kV JEM1010 (Jeol) with a CMOS 4x4 sensor (TemCam F416). **B.** RELION 2D averages of gp15. The C-terminal region is pointed with an arrow. **C.** C6 symmetrized gp15 3D reconstruction generated with RELION. The resolution is estimated to be 20 Å. **D.** A representative micrograph of gp16 obtained on a JEM1010 (Jeol) using a CMOS 4x4 sensor (TemCam F416). **E.** A representative micrograph of the gp15-gp16 complex obtained on a JEM1010 (Jeol) with a CMOS 4x4 sensor (TemCam F416). **F.** 2D average images of gp15-gp16. **G.** C6 symmetrized gp15-gp16 3D model at 22 Å resolution. The scale bar is 40 Å.

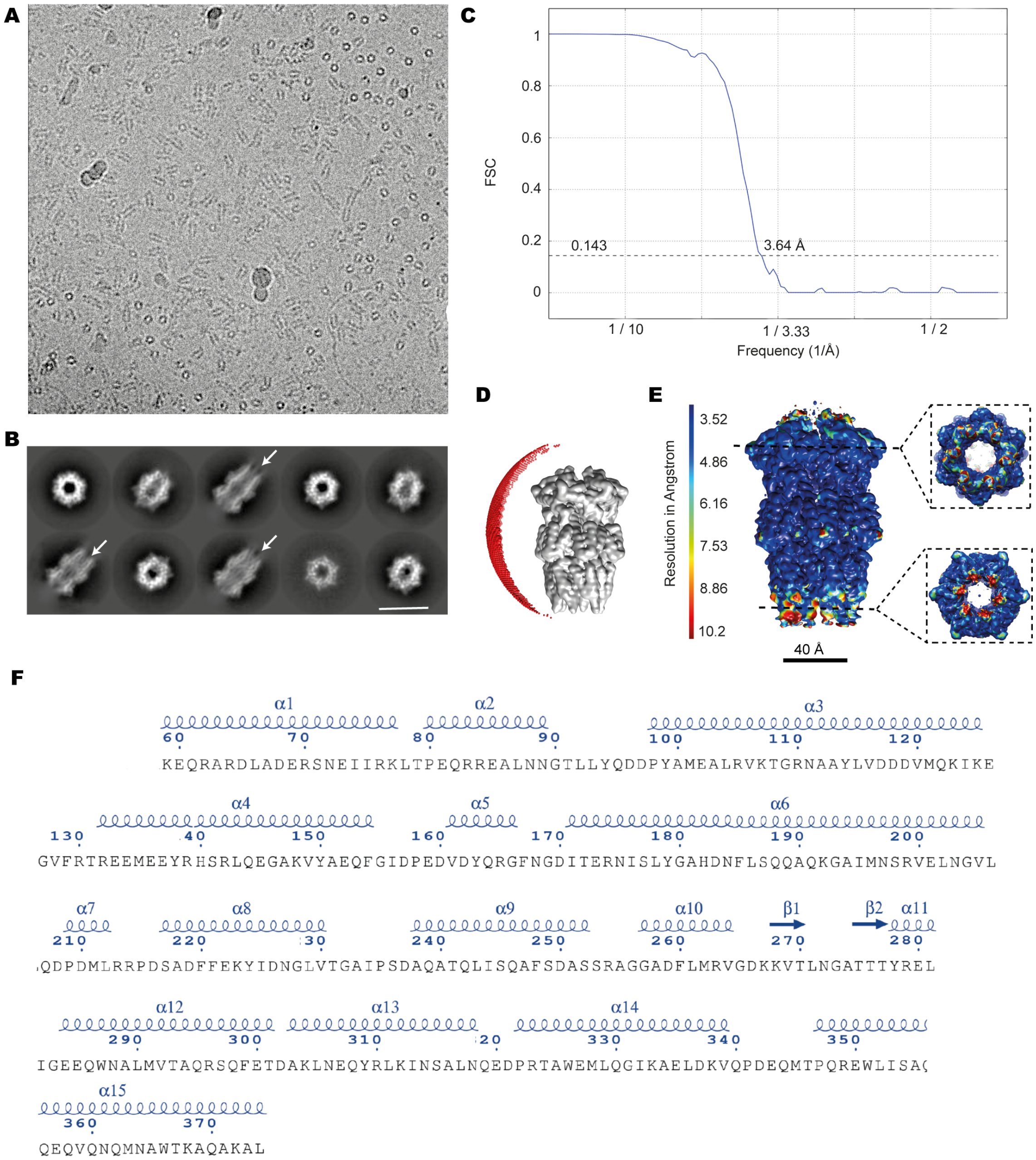

**Supplementary Figure 3. Cryo-EM analysis of bacteriophage T7 gp15. A.** A representative micrograph obtained on a 200kV Talos Arctica microscope (Thermo Fisher) with a Falcon III direct electron detector. **B.** 2D average images of gp15 (scale bar = 140 Å). The C-terminal region is pointed with an arrow. **C.** Gold-standard Fourier Shell Correlation (FSC) curve. The resolution was estimated at FSC = 0.143 (black dashed line) to 3.64 Å. **D.** 3D model of gp15 showing Euler angle distribution (red dots) of particles used during the refinement; a C6 symmetry was imposed. **E.** Side view of the gp15 3D model showing the local resolution estimation performed with MonoRes in angstroms (Å). Two insets show the resolution of the inner channel at two different axial planes. The highest resolution was 3.52 Å and the lowest was 10.2 Å. **F.** Secondary structure chart of the gp15 protein traced obtained with ENDscript 2.0 (1); α-helixes, β-sheets and residue numbers are labeled.

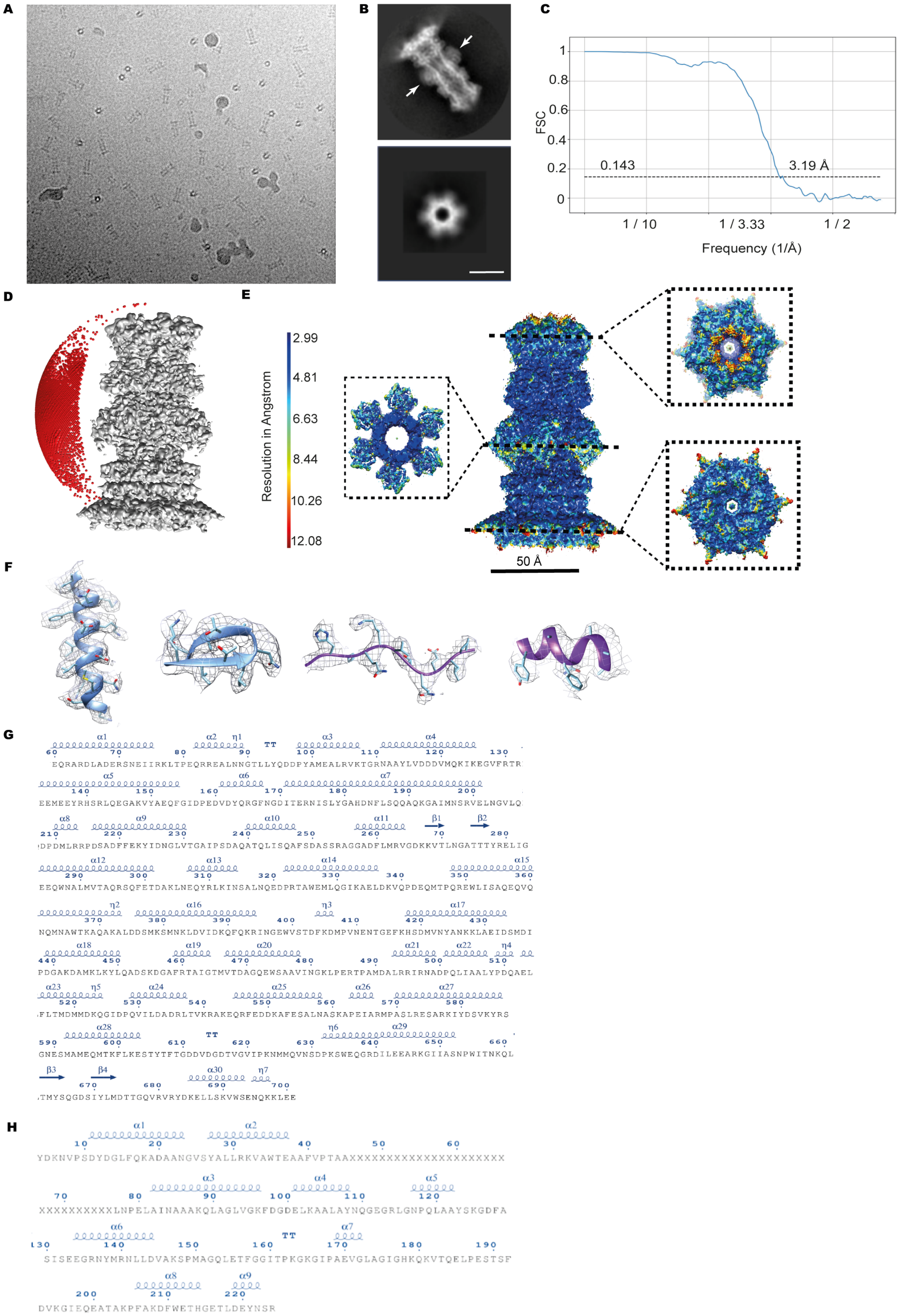

**Supplementary Figure 4. Cryo-EM analysis of the gp15-gp16 core complex. A.** A representative micrograph of gp15-gp16 core complex obtained on a 300kV Titan Krios microscope (FEI) equipped with K2 Summit (Gatan) direct electron detector. **B.** 2D classes of axial (left) and side view (right) of the gp15-gp16 complex; scale bar = 210 Å. **C.** Gold-standard Fourier Shell Correlation (FSC) curve indicating the resolution of the density map (light blue line). The resolution was estimated to be 3.19 Å using the FSC = 0.143 criteria (black dashed line). **D.** C6 symmetrized 3D model of gp15-gp16 showing the Euler angle distribution of the particles used in the refinement. **E.** Side view of the gp15-gp16 3D model showing the local resolution estimated with MonoRes in angstrom (Å). The insets show the resolution of the model inside the channel at three different axial planes. **F,** Extraction of the density from the 3D reconstruction corresponding to a α-helix and β-sheet from gp15 (blue) and loop and α-helix from gp16 (purple) showing the placement of the side chains. Notice that even though the chosen gp16 α-helix is found at the lower resolution region of the structure big side chains can be appreciated. **G,H.** Secondary structure chart of gp15 (g) and gp16 (h) traced in the complex obtained with ENDscript 2.0 software (1), α-helixes, β-sheets and residue number are labeled.

**Supplementary Table 1. Cryo-EM data collection, refinement and validation statistics.**

| | **gp15 tubular core protein (EMD-10911, PDB 6YSZ)** | **Gp15-gp16 core complex (EMD-10912, PDB 6YT5)** |
|---|---|---|
| **Data collection and processing** | | |
| Microscope | Talos Arctica | Titan Krios |
| Voltage (kV) | 200 | 300 |
| Camera | Falcon III | K2 Summit |
| Nominal magnification | 120000 | 47619 |
| Exposure time (s) | 35 | 8 |
| Frames per movie | 62 fractions<br>22 frames/fraction | 32 |
| Electron exposure (e–/Å$^2$) | 30 | 40 |
| Defocus range (μ m) | -1,4 to -2,4 | -1,0 to -2,8 |
| Pixel size (Å) | 0.855 | 1.047 |
| Symmetry imposed | C6 | C6 |
| Micrographs (no.) | 2470 | 5506 |
| Particle images (no.)[a] | 50980 (63829) | 72882 (274918) |
| Map resolution (Å)<br>FSC threshold | 3,64<br>0.143 | 3,19<br>0.143 |
| Map resolution range (Å) | 3.52 Å – 10.2 Å | 2.99 Å – 12.08 Å |
| **Refinement** | | |
| Initial model used | gp15 full-length (6YT5) | gp15 none (*ab-initio)*, gp16 (threading with 1SLY and 4CFO) |
| Map-sharpening B factor (Å$^2$) | n.a. | n.a. |
| **Model composition** | | |
| Nonhydrogen atoms | 15192 | 39078 |
| Protein residues[b] | 1896 | 5040 |
| **Protein** | | |
| FSC average | 0.89 | 0.90 |
| R factor | 0.31 | 0.29 |
| **R.m.s. deviations** | | |
| Bond lengths (Å) | 0.003 | 0.007 |
| Bond angles (°) | 0.541 | 0.614 |
| **Validation** | | |
| MolProbity score | 1.76 | 2.12 |
| Clash score | 10.46 | 13.10 |
| Rotamers outlayers (%) | 0 | 0.15 |
| **Ramachandran plot** | | |
| Favored (%) | 96.55 | 91.77 |
| Allowed (%) | 3.45 | 8.23 |
| Disallowed (%) | 0 | 0 |

[a] The initial number of images is shown in parentheses.
[b] Number of protein residues in each chain.

REFERENCES

1. X. Robert, P. Gouet, Deciphering key features in protein structures with the new ENDscript server. *Nucleic Acids Res* **42**, W320-324 (2014).
